# Comparative analysis of mandible morphology in four ant species with different foraging and nesting habits

**DOI:** 10.1101/2021.08.26.457866

**Authors:** Ritabrata Chowdhury, Neelkamal Rastogi

## Abstract

Mandibles of ants can be considered as one of the most vital tools for the survival and success of their colonies since these are extensively utilised for defence, nest maintenance and foraging activities. We hypothesised that mandibular design is strongly dependent on the respective ecological niche and foraging habit of an ant species. In the present study, we compared the external morphology and zinc content in the mandibles of four species of ants by using scanning electron microscopy (SEM) and energy dispersive X-ray spectroscopy (EDX). The ant mandible morphology varied significantly in accordance with their species-specific foraging and nesting strategies. The sickle-shaped mandibles of the strongly predaceous, *Oecophylla smaragdina* worker ants were characterised by a large number of pointed teeth which would be of immense utility for subduing the prey, while the shovel-shaped, highly sclerotized mandibles of *Cataglyphus longipedem* ants appear to be adaptations for the solitary scavenging habit and nest maintenance in arid habitats. The large-sized mandibles of *Camponotus compressus* ants and the stout mandibles of the predatory, *Tetraponera. rufonigra* forager ants, have apparently evolved for collection of sugary secretions by the former and for the solitary foraging and arboreal nesting habit of the latter. The mandibular zinc content was highest in *T. rufonigra* ants and the lowest in the mandibles of the sugar-loving *Cm. compressus* ants. The diversity in the arrangement of bristles and the type of mandibular concavities, have also evolved accordingly. Thus, this study might prove to be instrumental in evaluating the various physical mechanisms involved in the evolution of insect mandibles for their defined function.

## 1. Introduction

Mandibles are crucial tools for most insects (Blanke et al. 2017) and have acquired various structural features according to their respective functions. These are well-developed in most species of ants which are basically omnivorous, although many species are predominantly predaceous and/or scavengers or feed mainly on plant-derived sugary secretions (Agarwal and Rastogi 2008a, 2008b; Khan et al. 2017). Ants use their mandibles for an extensive range of activities and consequently, have evolved a spectacular diversity in their foraging strategies (Höldobbler and Wilson 1990). The worker ants use their mandibles primarily to manipulate food items of varying sizes, and even liquids (e.g. water or honeydew suspended as a droplet between the mandibles). Like other eusocial insects, ants often construct elaborate nests, using their mandibles to dig into the dirt or wood and to carry and dump the debris outside the nest. Attendant ant workers frequently move brood to other nest chambers using their mandibles and in some species, social carrying of workers by means of mandibles is also reported (Zhang et al. 2020).

In the present study, we have aimed to examine and compare the mandible morphology of four species of ants (Hymenoptera: Formicidae) from four different genera, and with vastly different foraging and nesting niches. While *Oecophylla smaragdina* (Fabricius), *Cataglyphis longipedem* ((Eichwald) (*Ct. longipedem*, hereafter) and *Camponotus compressus* (Fabricius) (*Cm. compressus*, hereafter) are members of subfamily Formicinae, *Tetraponera rufonigra* (Jerdon) belongs to subfamily Pseudomyrmecinae..

The Asian weaver ant, *O. smaragdina*, is characterised by a territorial organisation and dominance in arboreal tree mosaics (Dejean et al 2007). Cooperation among the worker castes is an important underlying feature of both the arboreal leaf nests constructed from leaves still attached to the trees and the foraging success of the ferociously predatory weaver ant colonies (Vanderplank 1960; Rastogi 2000, 2007). They are group foragers and being predaceous, exhibit extensive prey handling behaviour (Rastogi, 2000). Large insect prey and intruders are often spread-eagled by several workers and killed by prolonged stretching (Ledoux 1950). While *T. rufonigra* is a solitary foraging, predatory species, with arboreal nests usually established in holes previously bored by wood-boring insects in the tree trunks (Norasmah et al 2012), *Ct. longipedem*, is a ground-nesting desert ant species adapted to harsh conditions of arid and semi-arid areas of Central and South Asia (Bharti et al 2015, Khan et al. 2017). The worker ants are primarily scavengers, and forage individually on dead arthropods. They build permanent nests in the hard, arid soil and use their mandibles for carrying out most activities. However, the carpenter ant, *Cm. compressus*, is a ground-nesting, extrafloral nectar foraging species (Ekka and Rastogi 2019) which builds its long-lasting main nest in moist wood, especially at the base of trees (Ayyar 1935) and temporary satellite nests in moist soil (Kumari et al 2016). They tend plant-sap-sucking insects like aphids and treehoppers and preferentially visit plants with extrafloral nectaries and/or those harbouring honeydew excreting hemipterans (Kumari and Rastogi 2018). All foraging, brood care, and nest maintenance activities carried out by ants are done by means of their mandibles. Thus, examination of the mandible morphology of ant species with different foraging strategies can provide interesting insights regarding the structure-function relationships in ants.

The main objective of the present study was to compare the mandible morphology of the ant foragers (i.e. the minor caste ants) of each of the four species: *O. smaragdina, T. rufonigra, Ct. longipedem* and *Cm. compressus*, and to co-relate their mandible characteristics to their forging behaviour, adapted niche and ecology. Here, we have focused on the morphometric parameters, scanning electron microscopy and electron-dispersive spectroscopy to examine the variations in mandible morphology across the ant species with varied food preferences and foraging modes.

## 2. Material & methods

Minor caste worker ants of *O. smaragdina, T. rufonigra, Ct. longipedem* and *Cm. compressus* were collected by the hand picking method, (during 2019), from the Botanical Garden of Banaras Hindu University Campus, Varanasi (25.2677° N, 82.9913° E), Uttar Pradesh, India. Ants were stored in vials containing 75 % ethanol and brought to the laboratory. Since the locally occurring ant species have been under investigation by various researchers in our group since more than two decades, all have been already identified by experts from ZSI, Kolkata, India.

Both the left and right mandibles of the ant foragers were dissected out using a stereo-binocular microscope (Leica MZ6 Stereomicroscope, Leica AG, Germany). The samples were prepared for SEM analysis by fixing them in 2.5% glutaraldehyde solution, followed by dehydration and drying in a desiccator. The samples were then carefully mounted in pairs using carbon tape and the mandibles were sputter coated with a thin gold layer (Quorum Sputter Coater).

Scanning electron microscope (SEM, Carl Zeiss EVO 18 Research, Carl Zeiss AG, Germany) was used to image the samples with accelerating voltages between 5and 20kV. To quantify the elemental composition of mandibles, EDX detector in the SEM was used to scan the samples and the spectra collected using an accelerating voltage of 20 kV. Variations in zinc content and other metals in mandibles were determined by using point Energy dispersive X-ray spectroscopy (EDX) analysis and elemental maps. Processing of electron micrographs was done by using Image J software.

An analysis of variance (ANOVA), Kruskal-Wallis test and the Principal Component Analysis (PCA) were used to analyse the variations in the mandible morphometric data and also Zinc content (wt in %) of the mandibles, among the four ant species under study, by using PAST 4.04 (Hammer et al 2001). The number of principal components (PCs) was determined by considering eigenvalues higher than those generated by the broken-stick method.

## 3. Results

The results of SEM showed that the mandibles of *Ct. longipedem* were large and shovel-like, strongly sclerotized, bearing 6-7 teeth on the outer margins (Fig 1.e-f), whereas in *O. smaragdina*, the mandibles were rather narrow and elongated. Along the entire length of the cutting edge, a concave depression was found which gave a sickle-like aspect to the mandible*s* of *O. smaragdina*. The first or the distal tooth being greatly elongated as compared to the other teeth, the mandibles of the weaver ant workers were characterised by the presence of 11-13 teeth, which generated a serrated pattern along the outer margin (Fig 1.c-.d; Table 1.). The bristles were distributed throughout the mandibular surface in *Ct. longipedem* ants but in *O. smaragdina* foragers, these were located specifically along the margin, just behind the cutting edge (Fig. 1c-d). The mandibles of *Cm. compressus* foragers were large, well-sclerotized with 6 teeth in each, similar to those recorded in the mandibles of *Ct. longipedem* ants. Bristles were found scattered on the entire surface of the mandibles (Fig. 1.a-.b). However, in case of *T. rufonigra* foragers, the mandibles were found to be rather short, stout and rectangular shaped, each bearing 5 teeth and a large number of bristles, which were mostly concentrated along the inner margin, bordering the teeth while a conspicuous concavity was present on the ventral side of each mandible (Fig 1.g-.h).

**Fig.1.**
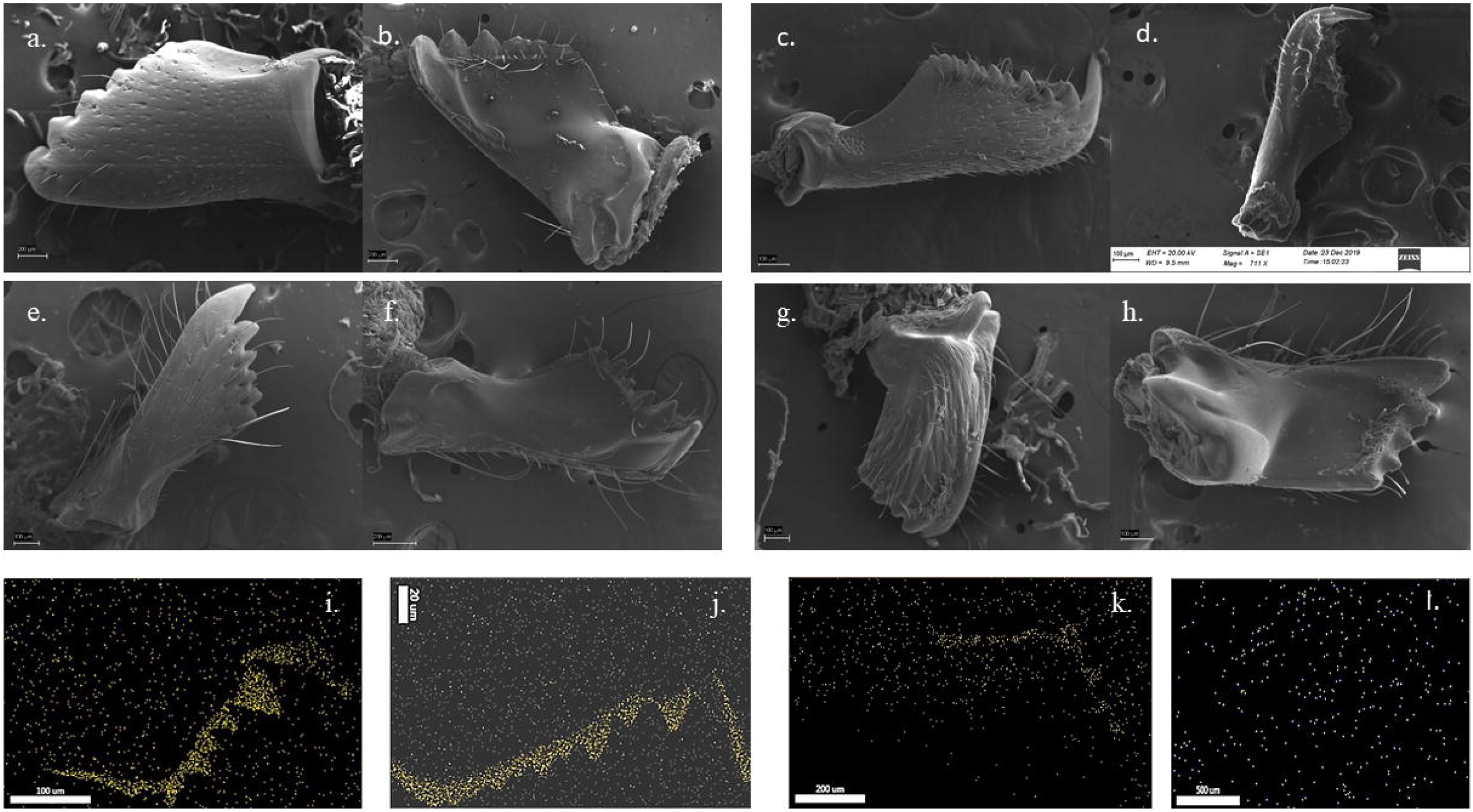
Scanning electron microscopy and electron dispersive X-ray spectroscopy analysis of mandibles in four ant species: **a** - **b** *Camponotus compressus* mandibles, in dorsal and ventral view, **c** - **d** *Oecophylla smaragdina* mandibles in ventral view. **e** - **f** *Cataglyphis longipedem* mandibles, in dorsal and ventral view. **g** - **h** *Tetraponera rufonigra* mandibles, in doral and ventral view; **i - l** EDX-elemental maps showing the distribution of zinc in the mandibles of *Ct. longipedem, O. smaragdina, T. rufonigra* and *Cm. compressus* respectively.

EDX was used to characterize the presence of elements, namely Zn, Mn, Fe, Cu and Na, at different spatial locations of the mandibles. Analyses of the mandibular surface showed peaks of zinc content, corresponding to the zinc located preferentially at the tip regions of the mandibles but not at the basal mandibular regions (Fig-2.a -d). The presence of zinc was found to be exclusive to the cutting regions of the mandibular apical regions whereas the neighbouring regions were devoid of zinc. This trend was further supported by results of the elemental maps (Fig-1 i-l). Other elements such as sodium, manganese (Mn) and iron (Fe) were also found in the elemental maps but due to their presence in extremely insignificant amounts, we have not considered them in the present report. Variations were found in the amount of zinc content (wt in %) present at the tips of the mandibles (Table 1.), with the highest amount being present in the mandibles of *T. rufonigra*, followed by that found in mandibles of *O. smaragdina, Ct. longipedem*, while the concentration was lowest in *Cm. compressus* mandibles. However, zinc content of *Cm. compressus* mandibles and also its distribution was considerably inconspicuous as compared to that recorded in the mandibles of other three ant species.

**Fig. 2.**
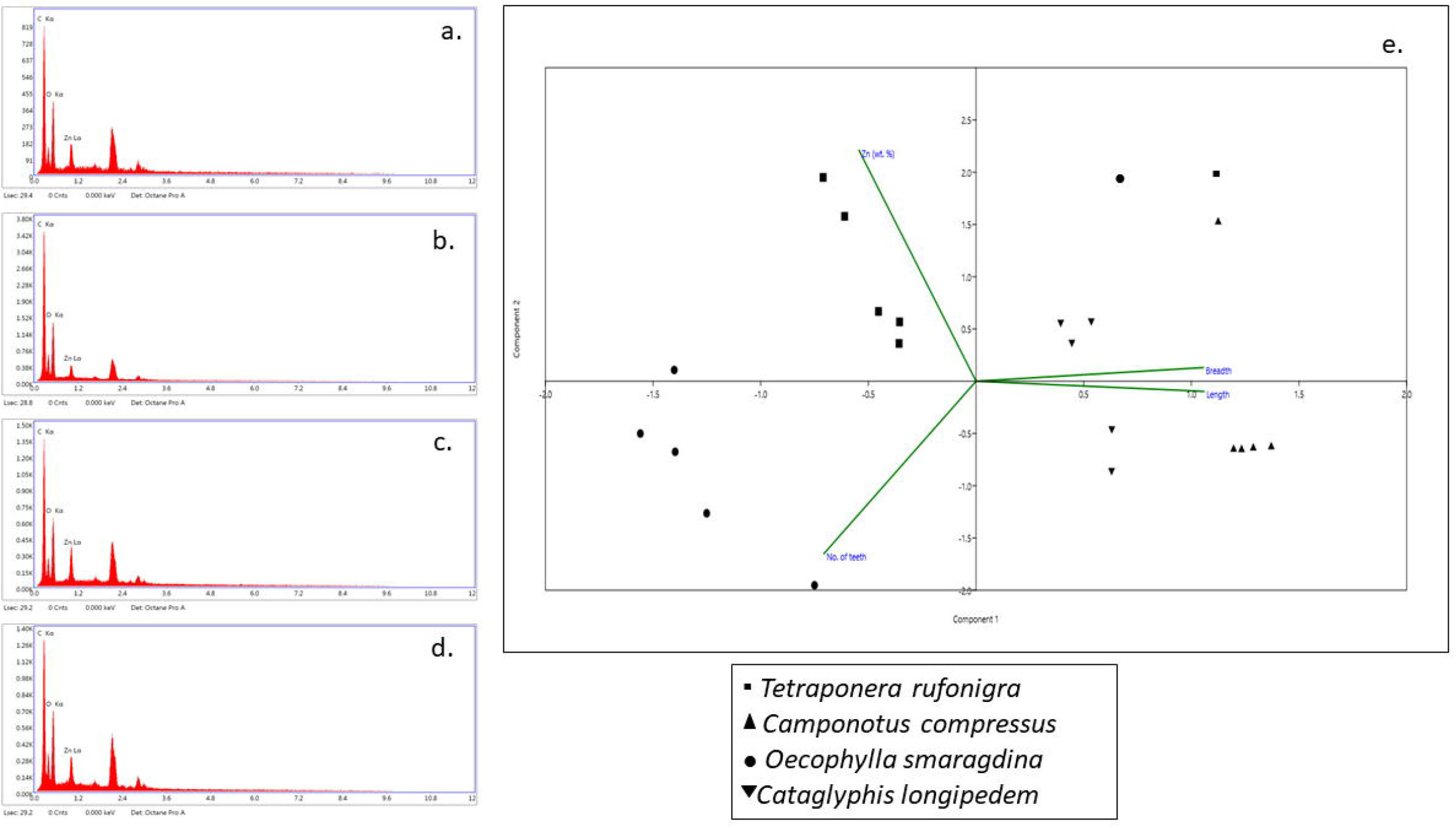
a-d EDX point plots and data analysis of the morphometric parameters of mandibles of the four ant species: **a**. *Camponotus compressus*, **b**. *Cataglyphis longipedem*, **c**. *Tetraponera rufonigra*, and d. *Oecophylla smaragdina*; **e**. Principal Component Analysis (PCA) to analyse the relationship between the mandible morphometric data and Zinc content (wt in %) of the mandibles, among the four ant species: *Camponotus compressus, Cataglyphis longipedem, Tetraponera rufonigra*, and *Oecophylla smaragdina*

Principal component analysis using morphometric characteristics and zinc content (Fig. 2.e) produced two axes that together explained the total variance. Significant differences were found in the mandibular features of the four ant species (p<0.05).

## 4. Discussion

Our results revealed a well-marked relationship, in each case, between the characteristic foraging strategy and the morphometric features, Zn content and its distribution in the mandibles of each of the four ant species, under study. The shovel-shaped mandibles of the scavenging, *Ct. longipedem* forager ants, appear to be well-adapted for simple retrieval of dead arthropod prey and for nest construction in the harsh arid ground which it usually inhabits (Khan et al. 2017). It would also facilitate the carrying of the colony individuals during nest shifting periods as reported in an earlier study (Lenoir et al. 2009). The worker ants of this ground–nesting, desert species, being solitary foragers, are single-prey loaders and scavengers which search and retrieve dead arthropods (Wehner et al. 1983; Cerdá 1988). The highly aggressive foragers of the arboreal colonies of *O. smaragdina* are characterised by group handling and retrieval of large prey and the unique shape and structure of their mandibles is suitably adapted for cooperative defence and prey capture. The elongated distal tooth and the concavity of the sickle shaped mandibles would greatly help in obtaining a firm grip of a struggling prey during cooperative prey capture and for its transport to the nest as shown in earlier studies (Polidori 2011; Fowler et al, 1986). Evidently, the large number of mandibular teeth (ranging from 11 to 13) is an effective adaptation for cutting large-sized prey into pieces. These adaptations in the mandibles of *Oecophylla* ants provide support to earlier studies of their foraging success on large prey (Rastogi 2000), in defending three-dimensional territories (Dejean et al. 2007; Rastogi 2007), in nest defence (Hölldobler and Wilson 1990) and also in cooperative construction of leaf nests (Dejean et al. 2007). The mandibular configuration of tree ants appears to be similar to that found in the mandibles of the leaf-cutting ant (Fowler et al. 1986). The shorter mandibles of *T. rufonigra* ants were markedly different from those of *O. smaragdina*, and appear to be adapted to enable the individually foraging workers to efficiently grip and carry the prey to their nest reportedly located at great heights from the ground (Norasmah et al. 2012).

The broad mandibles of the sugar-loving, *Cm. compressus* ant foragers (Kumari and Rastogi 2018) with a few (4-5) short and conical teeth, showed some morphological similarity to *Ct. longipedem* mandibles. The short mandibles of the carpenter ants have apparently evolved for foraging directly and/or indirectly on plant-derived liquid resources, such as extrafloral nectar and honeydew excreted by insect symbionts (Agarwal and Rastogi 2010). Carpenter ants documented to be far less aggressive than ecologically dominant ants such as *O. smaragdina* and *Pheidole latinoda* (Rastogi 2007) exhibit only moderate level of predatory impact on insect prey (Agarwal and Rastogi 2008a). Apart from using their mandibles for feeding on the liquid carbohydrate sources they also use it for brood care, for digging their nests in the soft soil/wood and for group foraging activities as reported in an earlier study (Ayyar 1935).

The differences in mandible morphology of the four species are clearly structural modifications which have evolved due to their diversified foraging habits and distinctly varied ecological niches. The long, slender, serrated mandibles with elongated distal teeth of *O. smaragdina* are adaptations necessary for its arboreal habitat and predominantly predatory habit, whereas the large, broad and stout mandibles with lesser teeth recorded in *Ct. longipedem* and *Cm. compressus*, respectively, indicate their scavenging and liquid food collecting habits, both of which do not involve active resistance from a struggling prey as is found in the case of the predatory weaver ant species. Further, the distribution of the bristles is of importance in this respect, as they seem to aid the ants in better gripping of the foraged items (Zhang et al. 2020) and this presumably explains the larger number of bristles recorded along the cutting margin of mandibles in arboreal, predatory ant species.

Interestingly, the varied content of zinc at mandible apical parts can also be linked to the variations in the feeding behaviour and habits of the respective ant species. (Fig 1). Presence of zinc has been found to contribute towards greater hardness to the mandibular edges (Cribb et al. 2008) and thus, higher amounts of zinc may indicate higher degree of hardness (Schofield et al. 2002). Our results also show that the Zn content was highest in case of *T. rufonigra*, whose workers are arboreal, solitary foragers, capable of individually carrying the caught prey up to large heights. Similarly, the higher concentration of Zn in *O. smaragdina* worker mandibles may be co-related to their actively predatory, arboreal habit, wherein large prey are subdued, cut into small pieces and carried up to their leaf nests by workers using their mandibles. Surprisingly, *Ct. longipedem* mandibles showed Zn content similar to that found in the *Oecophylla* ants. Earlier studies have reported a high preference of *Ct. longipedem* colonies for heavy metal polluted areas in comparison to un-polluted and/or less polluted areas (Khan et al. 2017). This preference is evidently related to their solitary, foraging behaviour and nesting activities in a xeric and rocky terrain (Khan et al. 2017), which necessitates the presence of strong mandibles, an adaptation which involves the deposition of heavy metals, such as zinc. Lastly, the mandibles of the sugar-loving, *Cm. compressus* ants exhibited the lowest Zn concentration, as it is primarily a nectar-feeding, species and the mandibles are used by workers for tasks such as grooming, tending the brood and for digging nests in soft soil/wood. Even though ant mandibles might have similar microstructures as shown in a recent study (Barlow et al. 2020), their morphology and design can be co-related to their respective niches and foraging habits.

To conclude the present study provides an insight into the significant morphological differences in the mandibles (in terms of its shape, size, presence and distribution of bristles, the presence and type of concavities and the zinc content) of the worker ants of the four different species to shed some light on the mechanisms involved in their diversification. However it is not without a few caveats. Firstly, absolute quantification of zinc has not been done and future studies need to focus on this aspect. Secondly, more intensive biomechanical analysis is required to delve deeper into the structure-function relationships. This might help us to develop a better understanding of the mechanisms involved in the evolution and biomechanics of insect mandibles.

## Declarations

### Author’s contributions

All authors had full access to all the data in the study and take responsibility for the integrity of the data and the accuracy of data analysis.

## Acknowledgement

We would like to acknowledge Prof. N.V. Chalapathi Rao, Department of Geology, Institute of Science, Banaras Hindu University, Varanasi, for kindly allowing us to utilise the SEM-EDS facility at his lab.

## References

1. Agarwal VM, Rastogi N (2008) Role of floral repellents in the regulation of flower visits of extrafloral nectary-visiting ants in an Indian crop plant. Ecol Entomol 33:59–65.

2. Agarwal VM, Rastogi N (2008) Deterrent effect of a guild of extrafloral nectary-visiting ant species on Raphidopalpa foveicollis, a major insect pest of sponge gourd, Luffa cylindrica. Entomol Exp Appl 128:303–311.

3. Agarwal VM, Rastogi N (2010) Ants as dominant insect visitors of the extrafloral nectaries of sponge gourd plant, Luffa cylindrica (L.) (Cucurbitaceae). Asian Myrmecol 3:45–54.

4. Ayyar PK (1935) The biology and economic status of the common black ant of south India-Camponotus (Tanaemyrmex) compressus, Latr. Bull Entomol Res 26(4):575–58.

5. Barlow MM, Bicknell RDC, Andrew NR (2020) Cuticular microstructure of Australian ant mandibles confirms common appendage construction. Acta Zool 101:260–270.

6. Bharti H, Guénard B, Bharti M, Economo EP (2016) An updated checklist of the ants of India with their specific distributions in Indian states (Hymenoptera, Formicidae). ZooKeys 551:1–83.

7. Blanke A, Watson PJ, Holbrey R, Fagan MJ (2017) Computational biomechanics changes our view on insect head evolution. Proc R Soc B 284:20162412.

8. Cribb BW, Stewart A, Huang H, Truss R, Noller B, Rasch R, Zalucki MP (2008) Insect mandibles— comparative mechanical properties and links with metal incorporation. Sci Nat 95:17–23.

9. Dejean A, Corbara B, Orivel J, Leponce M (2007) Rainforest canopy ants: the implications of territoriality and predatory behaviour. Funct Ecosyst Communit 1:105–120.

10. Ekka PA, Rastogi N (2019) A single lycaenid caterpillar gets an ant-constructed shelter and uninterrupted ant attendance. Entomol Exp Appl 167:1012–1019.

11. Fowler HG, Pereira-da-Silva V, Forti LC, Saes NB (1986) Population dynamics of leaf-cutting ants: a brief review. pp. 123–145. In: Lofgren, C. S.; Vander Meer, R. K. (Eds.). Fire ants and leaf cutting ants: Biology and management. West-View Press, Boulder. Colorado. USA.

12. Hammer Ø, Harper DAT, Ryan PD (2001) PAST: Paleontological statistics software package for education and data analysis. Palaeont Electron 4:9.

13. Hillerton JE, Robertson B, Vincent JF (1984) The presence of zinc or manganese as the predominant metal in the mandibles of adult, stored-product beetles. J Stored Prod Res 20:133–137.

14. Hingston, RWG (1927) The habits of Oecophylla smaragdina. Proc Ent Soc 2:90–94.

15. Hölldobler B (1984) Konkurrenzverhalten und Territorialitat in Ameisenpopulationen.In: Eisner T.et al. (Eds) Chemische Okologie, Territorialitat, Gegenseitige Verstandigung (Information processing in animals, vol. 3). New York: Gustav Fisher Verlag, pp. 25–70.

16. Hölldobler B, Wilson EO (1990) The Ants. Belknap Press of Harvard University, Cambridge, MA.

17. Khan SR, Singh SK, Rastogi N (2017) Heavy metal accumulation and ecosystem engineering by two common mine site-nesting ant species: implications for pollution-level assessment and bioremediation of coal mine soil. Environ Monit Assess 189:195.

18. Kumari S, Singh H, Rastogi N (2016) Influence of the sugar-loving ant, Camponotus compressus (Fabricius, 1787) on soil physico-chemical characteristics. Halteres 7:163–174.

19. Kumari S, Rastogi N (2018) Can a common and abundant plant-visiting ant species serve as a model for nine Sympatric ant-mimicking arthropod species? Curr Sci 114 (10):2189–2192.

20. Kundanati L, Chahare NR, Jaddivada, S, Karkisaval AG, Sridhar R, Pugno NM, Gundiah N (2020) Cutting mechanics of wood by beetle larval mandibles. J Mech Behav Biomed. Mater 112:104027.

21. Ledoux A (1950) Recherches sur la biologie de la fourmifileuse (Oecophylla longinoda Latr). Ann Sci Nat Zool 12:313–461.

22. Lenoir A, Aron S, Cerdá X, Hefetz A (2009) Cataglyphis desert ants: a good model for evolutionary biology in Darwin’s anniversary year - A review. Isr J Entomol 39:1–32.

23. Norasmah B, Chin YJ, Abu Hassan A (2012) A preliminary study on the diurnal foraging activity and nutrient preferences of Tetraponera rufonigra (Hymenoptera: Formicidae) in Pulau Pinang, Malaysia. Malays Appl Biol 41:51–54.

24. Polidori C (2011) Predation in the Hymenoptera: An Evolutionary Perspective. Transworld Research Network, Trivandrum, Kerala.

25. Rastogi N (2000) Prey concealment and spatiotemporal patrolling behaviour of the Indian tree ant Oecophylla smaragdina (Fabricius). Insectes soc 47:92–93.

26. Rastogi N (2007) Seasonal pattern in the territorial dynamics of the arboreal ant (Hymenoptera: Formicidae). J Bombay Nat Hist Soc 104(1).

27. Schofield RM, Lefevre HW (1992) PIXE-STIM microtomography: zinc and manganese concentrations in a scorpion stinger. Nucl Instrum Methods Phys Res 72:104–110.

28. Schofield RM, Nesson MH, Richardson KA (2002) Tooth hardness increases with zinc-content in mandibles of young adult leaf-cutter ants. Sci Nat 89:579–583.

29. Vanderplank FL (1960) The bionomics and ecology of the red tree ant, Oecophylla sp., and its relationship to the coconut bug Pseudotheraptus wayi Brown (Coreidae). J Anim Ecol 29:15–33.

30. Wehner R, Meier C, Zollikofer C (2004) The ontogeny of foragwehaviour in desert ants, Cataglyphis bicolor. Ecol Entom 29:240–250.

31. Zhang W, Li M, Zheng G, Guan Z, Wu J, Wu Z (2020) Multifunctional mandibles of ants: Variation in gripping behavior facilitated by specific microstructures and kinematics. J Insect Physiol 120:103993.

