## Supplementary material for "Comparative analysis of mandible morphology in four ant species with different foraging and nesting habits": Table-1

**Table 1**. Morphometric characteristics (mean±SEM) and average Zn content in mandibles of the four ant species (n=5 for each species).

| **Species** |  |  | **Length (µm)** | **Breadth (µm)** | **No. of teeth** | **Avg. Zn wt. (%)** |
| --- | --- | --- | --- | --- | --- | --- |
| **Tetraponera rufonigra** |  |  | 1025.41±10.60 | 439.09±4.31 | 5.00±0.00 | 11.77 |
| **Cataglyphis longipedem** |  |  | 1827.69±20.55 | 734.01±20.84 | 6.60±0.21 | 6.55 |
| **Camponotus compressus** |  |  | 2075.91±17.89 | 1007.50±45.73 | 6.00±0.00 | 3.44 |
| **Oecophylla smaragdina** |  |  | 928.194±67.56 | 330.62±17.70 | 12.00±0.28 | 8.79 |
